# COMPARATIVE GENOMIC ANALYSIS OF CORE AND ACCESSORY GENES IN RUST FUNGI REVEALS PATHOGENICITY-ASSOCIATED GENE FAMILIES IN *Phakopsora pachyrhizi*

**DOI:** 10.64898/2026.07.03.736376

**Authors:** Vinicius Delgado da Rocha, Liliane Santana Oliveira, Francismar Corrêa Marcelino-Guimarães

## Abstract

Accessory genes are thought to contribute to fungal adaptation and pathogenicity by modulating host immunity, while core genes play crucial roles in maintaining fundamental biological processes. Rust fungi (order Pucciniales) are obligate biotrophic plant-pathogens and infect economically relevant crops. Here, we characterize core and accessory gene repertoires across rust fungi, with a particular focus on *Phakopsora pachyrhizi*, the causal agent of Asian soybean rust. Across Pucciniales genomes, accessory genes represented the largest fraction of gene content (∼44.6% on average), whereas core genes accounted for a smaller proportion (∼18–35%). Notably, variations in accessory gene content among rust fungi are perhaps attributed to lineage-specific gene expansions and losses. Core gene content was positively correlated with total gene number across Pucciniales genomes, suggesting retention after gene duplication events, consistent with their essential biological functions. Among *P. pachyrhizi* genes expressed during soybean infection, core effectors were associated with cysteine-rich proteins, pectin-degrading enzymes, and SPFH/Band 7 family, while accessory effectors included phosphatidylethanolamine-binding proteins, trehalose phosphatases, and CFEM domain-containing proteins. The *in-plant* induced core and accessory genes in *P. pachyrhizi* also comprised multiple families of CAZymes (GH5/GH7 cellulases, CE5 cutinases, CE8 pectinesterases, CE4/GH18 chitin-modifying enzymes); proteases (aspartyl proteases, serine carboxypeptidases, alpha/beta hydrolases); transporters (amino acid permeases, ferric reductase-like transmembrane proteins, and OPT oligopeptide transporter), and transcription factors (bZIP, GATA zinc finger, STE-like, and homeobox KN). Our study highlights that core and accessory gene families have shaped *P. pachyrhizi*-soybean interactions, identifying promising targets for functional studies aimed at elucidating host-adaptation mechanisms in rust fungi.

## INTRODUCTION

Core genes are defined as a conserved set of genes shared among all members of a taxonomically related group (Jiao et al., 2018). The functional roles of core genes are typically associated with essential biological processes, such as cellular metabolism and reproduction, thereby promoting the organism’s survival (Croll & McDonald, 2012). Generally, core genes tend to experience a lower incidence of recombination and horizontal gene transfer, and exhibit highly conserved nucleotide sequences at the intra-specific level, with relatively reduced mutation rates due to constrained purifying selection (Sarkar & Guttman, 2004; Croll & McDonald, 2012).

While core genes are broadly conserved across species, accessory genes are present exclusively in a specific subset of species or strains (Badet & Croll, 2020). Genomic accessory regions often harbor high densities of transposable elements (TEs), display significant sequence polymorphisms, and can be acquired via horizontal gene transfer between isolates (Witte et al., 2021). Accessory genes are believed to confer adaptive functions, particularly in the context of complex molecular interactions between microbial plant-pathogens and their hosts (Croll & McDonald, 2012). In many fungal plant-pathogens, accessory genomic elements encode virulence-related factors, including effectors and host-selective toxins, which enable the manipulation of host immunity (Bertazzoni et al., 2018). For example, genes localized on accessory chromosomes are required for the biosynthesis of host-specific AM-toxins in the plant-pathogenic fungus *Alternaria alternata* (Witte et al., 2021). The importance of accessory genes in mediating virulence and host adaptation has been demonstrated experimentally. For instance, a recent functional study showed that three accessory genes (GPCGs 1, 16, and 17) in the polyphagous fungus *Colletotrichum fructicola* are required for lesion formation on apple leaves (Liang et al., 2024). Similarly, knockout mutants of two accessory genes (SIX1a and SIX4) in the necrotrophic fungal pathogen *Fusarium oxysporum* f. sp. *cubense* exhibited a substantial reduction in disease severity on banana rhizomes (Zhang et al., 2024). Additionally, a set of evidence also suggests that the acquisition of specific accessory genes in *F. oxysporum* f. sp. cubense tropical race 4 (TR4) may enhance fungal nitric oxide production, thereby suppressing host defenses and facilitating infection (Zhang et al., 2024).

The core and accessory gene content remains largely unknown within the order Pucciniales (rust fungi). Pucciniales constitutes one of the largest groups of plant-pathogenic fungi, comprising roughly 8,000 recognized species (Henningsen et al., 2025). This order includes obligate biotrophic fungi capable of infecting agriculturally important crops such as corn, coffee, oat, poplar, soybean, and wheat (Henningsen et al., 2025). The coevolution between rust fungi and their hosts is highly complex, and host shifts likely served as a primary driver of diversification (McTaggart et al., 2015). While the vegetative cycle of Pucciniales involves the production of dikaryotic urediniospores with independent haploid nuclei (N+N), its complete macrocyclic life cycle comprises up to five spore types and can occur either on a single host species or alternately between two distinct host plant species (Lorrain et al., 2019). However, the life cycle of many rust fungi remains poorly understood across several species, including *Phakopsora pachyrhizi*, for which only the asexual urediniospore and teliospores stages have been observed (Bromfield et al., 1994). Recent advances in long-read sequencing, Hi-C technologies, and robust bioinformatic tools have enabled the generation of haplotype-phased and chromosome-level genome assemblies for rust fungi (Henningsen et al., 2025). The haploid genome size of rust fungi ranges from approximately 75 Mb to 1 Gb and is positively correlated with TE content (Henningsen et al., 2025). Notably, TE abundance shows substantial interspecific variation in Pucciniales, ranging from ∼ 47% in *Melampsora larici-populina* to ∼ 92% in *P. pachyrhizi* (Corre et al., 2025).

One destructive species of rust fungus is *P. pachyrhizi*, which can infect approximately 150 leguminous species (Hartman et al., 2011). *P. pachyrhizi* is responsible for Asian Soybean Rust (ASR) disease, causing severe premature defoliation (Hossain et al., 2024). The occurrence of ASR has been reported worldwide, covering Africa, Asia, Australia, the United States, and South America (Hossain et al., 2024). Notably, ASR can have a high impact on soybean production, with yield losses of up to 90% in cases of insufficient management (Godoy et al., 2016). The development of effective management strategies for ASR is difficult owing to the presence of alternative hosts that serve as inoculum reservoirs, long-distance spore dispersal, and fungicide insensitivity among fungal populations (Godoy et al., 2016; Chicowski et al., 2024). Seven soybean rust resistance loci, namely *Rpps*1-7, have been identified on the soybean genome, but no commercial soybean cultivars currently provide durable resistance (Chicowski et al., 2024).

Despite the major agronomic importance of *P. pachyrhizi*, current knowledge of its virulence factors remains limited, with only a few functional studies on effector proteins (De Carvalho et al., 2017; Qi et al., 2018; Bueno et al., 2022; Castanho et al., 2024). Under laboratory conditions, *P. pachyrhizi* cultivation is challenging because of its obligatory biotrophic lifestyle, which requires living host tissue for growth (Twizeyimana et al., 2010). This biological constraint hinders the manipulation of *P. pachyrhizi* samples to conduct comprehensive studies on pathogen-host molecular interactions.

Currently, three *de novo* genome assemblies of *P. pachyrhizi* are available, reaching a size of up to 1,25 Gb (Gupta et al., 2023). The obtained genomes of *P. pachyrhizi* represent three distinct isolates (UFV02, K8108, and MT2006) collected in South America. These isolates belong to a widespread evolutionary lineage and exhibit moderate-to-high virulence on soybean genotypes carrying *Rpp* loci (Rocha et al., 2025). The available genome-wide data from *P. pachyrhizi* provide new perspectives to explore the evolution of core and accessory genes in the pathogen. Investigating these gene categories offers a key step toward understanding the molecular mechanisms underlying pathogenicity in rust fungi, such as *P. pachyrhizi*.

This study characterizes core and accessory, and singleton components in Pucciniales species, focusing on *P*. *pachyrhizi*. We specifically aimed to address the following questions: (1) What are the respective sizes of the core, accessory, and singleton gene repertoires within Pucciniales genomes? (2) What are the predicted functional roles of core, accessory, and singleton genes from *P. pachyrhizi*? (3) What core and accessory gene families from *P. pachyrhizi* are expressed during soybean infection? (4) What are the degrees of genetic diversity within putative core and accessory effector orthogroups in *P. pachyrhizi*? To answer these questions, we applied comparative genomic approaches to analyze publicly available genomic data from 12 phylogenetically diverse Pucciniales species, including three independent genome assemblies for *P. pachyrhizi*. Using an orthology inference method, we identified the core, accessory, and singleton repertoires across Pucciniales. From previous gene annotations analyses for *P. pachyrhizi*, we examined the molecular functions and enriched GO/KOG functional categories among core, accessory, and singleton genes in the pathogen. Finally, we retrieved transcriptome profiling data from *P. pachyrhizi* to evaluate the expression patterns of core and accessory genes during host-pathogen interactions. Our research provides evolutionary insights into the diversification of core and accessory genes across Pucciniales order and contributes to the identification of genes involved in *P. pachyrhizi* pathogenicity.

## MATERIAL AND METHODS

### Genomic data acquisition

This study utilized a total of 12 Pucciniales species obtained from geographically distant countries and collected from different host crops (Supplementary Table S1). For six species (*Cronartium quercuum*, *Melampsora allii-populina*, *Melampsora americana*, *Melampsora larici-populina*, *Melampsora medusae*, and *P. pachyrhizi*), we retrieved complete datasets of non-redundant protein sequences from curated gene catalogues available on the JGI MycoCosm database (Grigoriev et al., 2014). For five additional species (*Puccinia coronata*, *Puccinia graminis*, *Puccinia polysora*, *Puccinia striiformis*, and *Puccinia triticina* ), we compiled predicted complete genomes and proteomes through public repositories: NCBI databases (https://www.ncbi.nlm.nih.gov/), CSIRO Data Access Portal (https://data.csiro.au/), and GitHub platform (https://github.com/jimie0311/Puccinia-polysora-genome). We also included proteome data from *Austropuccinia psidii*, which was generously provided by Dr. Peri Tobias (University of Sidney). The quality and completeness of genome assemblies and proteomes were assessed using BUSCO v6 (Simão et al., 2015) with the pucciniomycetes_odb12 dataset and default parameters. The BUSCO completeness scores are reported in Supplementary Table S1.

To remove redundant protein sequences from the previously obtained whole-proteome dataset, we applied CD-HIT software (Li & Godzik, 2006) with thresholds of ≥95% identity and ≥95% coverage. After filtering, we compiled a final dataset consisting exclusively of non-redundant protein sequences from all 12 Pucciniales species. This dataset, hereafter referred to as Puc12 dataset, was used in downstream analyses.

### Orthogroup prediction and phylogenomic analysis

Clusters of orthologous protein sequences (orthogroups) were inferred from the Puc12 dataset using OrthoFinder v2.5.5 (Emms & Kelly, 2015) with an inflation parameter of 1.5.

For an in-depth phylogenomic reconstruction across Pucciniales species, we utilized a set of 98 single-copy orthologous proteins previously identified by OrthoFinder. This set of protein sequences was aligned with MAFFT v7.453 (Katoh & Standley, 2013) using the L-INS-i algorithm. The resulting sequence alignment was trimmed using the TrimAL software (Capella-Gutierrez et al., 2009) with parameter automated1. Subsequently, a maximum likelihood (ML) phylogenomic analysis was conducted in IQ-TREE software v1.6.12 (Nguyen et al., 2015), with ten independent runs and 1,000 ultrafast bootstrap replicates. The bootstrap convergence criterion was 0.99 and followed the previous standards described by Hoang et al., (2018). The best partition scheme and the best-fitting evolutionary models were selected by the ModelFinder2 (Kalyaanamoorthy et al., 2017) as implemented in IQ-TREE. The consensus phylogenomic tree was edited using FigTree v1.2.4 (http://tree.bio.ed.ac.uk/software/figtree/).

### Identification of core, accessory, and singleton genes

The orthogroups predicted by OrthoFinder, along with their corresponding gene members, were classified into three categories: core, accessory, and singleton. The term “core” was used to indicate orthogroups and their associated genes that were present in all 12 Pucciniales species. The term “accessory” corresponded to orthogroups/genes found in more than one species but not shared across all species. The term “singleton” referred to orthogroups/genes that were exclusively found in a single genome from one species.

To assess the degree of similarity in repertoires of core and accessory genes among Pucciniales species, we conducted two separate Principal Component Analyses (PCA), utilizing the prcomp function in R 4.4.0 (https://www.r-project.org/). The first PCA analyzed the number of core genes per orthogroup across species, while the second PCA focused on the number of accessory genes per orthogroup. Pearson’s correlation (r) coefficients were estimated between the number of core and accessory genes and the total number of protein-coding genes in each Pucciniales genome, using R software.

### Functional annotation and enrichment analyses

To gain insights into potential functional roles of core, accessory, and singleton genes, we specifically examined three *P. pachyrhizi* isolates (UFV02, K8108, and MT2006). The gene functional annotations for each *P. pachyrhizi* isolate were recovered from previously published analyses (Gupta et al.2023; https://mycocosm.jgi.doe.gov/Phakopsora). We then explored how key functional categories, including CAZymes, proteases, transcription factors, transporters, secreted proteins, effector genes, and secondary metabolites, were distributed into core, accessory, and singleton repertoires from *P. pachyrhizi*.

Finally, we investigated whether specific molecular functions were significantly enriched within each gene category (core, accessory, and singleton) in *P. pachyrhizi*. For each of the three isolates (UFV02, K8108, and MT2006), we performed two independent functional enrichment analyses using Gene Ontology (GO) and Eukaryotic Orthologous Groups (KOG) annotations derived from Gupta et al., (2023). GO-based enrichment was carried out with the topGO R package (Alexa et al., 2023), whereas KOG-based enrichment was performed using the FunFEA R package (Charest et al., 2025). Each analysis consisted of three independent comparisons, in which one gene category was compared against the other two.

### Expression profiles of core and accessory genes from *P. pachyrhizi* during soybean rust

Expression levels of core and accessory genes from *P. pachyrhizi* were assessed during urediniospore germination and throughout a time course of soybean infection (10–72 hours post-inoculation). To achieve this, we retrieved Log2 Fold-Change (Log2 FC) data from a comprehensive set of differentially expressed genes (DEGs) identified in UFV02 and K8108 isolates over soybean-host interactions. The DEGs and corresponding Log2 FC values were obtained from the RNA-seq transcriptome study conducted by Gupta et al., (2023). Each DEG was classified as either core or accessory based on our prior analyses. To ensure accuracy in gene expression analyses, we retained only ‘top DEGs’ defined as genes exhibiting a Log2 FC ≥ 3 at least one expression time point in each isolate. Our final dataset included the top DEGs categorized into two groups: core and common accessories belonging to orthogroups shared between the UFV02 and K8108 isolates. This strategy allowed that the analyzed DEGs were highly expressed and shared across the two isolates. The pheatmap R package (https://cran.r-project.org/web/packages/pheatmap/index.html) was used to construct heatmaps representing expression patterns.

### Genetic diversity of core and accessory effectors from *P. pachyrhizi*

To investigate molecular polymorphism among putative core and accessory effectors from *P. pachyrhizi*, we analyzed orthogroups containing the top DEGs that were classified as core and common accessory effectors according to our preceding analyses. Diversity analyses were performed on distinct datasets comprising coding-DNA sequences (CDSs) from three *P. pachyrhizi* isolates: UFV02, K8108, and MT2006. Each dataset represented an orthogroup containing at least one top DEG identified as either a core effector or a common accessory effector. Multiple sequence alignments for each dataset were performed in MAFFT v7.453 (Katoh & Standley, 2013) using the L-INS-i algorithm. The DNAsp software v.6 (Rozas et al., 2017) calculated for each dataset the following diversity measures: number of segregating sites (S), nucleotide diversity (π), and Watterson’s theta per site (θ_w_).

## RESULTS

### Ortholog-based phylogenomic analysis in Pucciniales

By analyzing 98 single-copy orthologous protein sequences, we generated an ML phylogenomic tree that reconstructed relationships among 12 Pucciniales species (Figure 1A). The tree exhibited strong statistical support for all main nodes, with bootstrap values of 100. The phylogenomic structure of the tree revealed two major clades within Pucciniales. The first clade displayed higher species diversity and included members from three families: Pucciniaceae (*P. coronata*, *P. graminis*, *P. polysora*, *P. striiformis*, and *P. triticina*), Sphaerophragmiaceae (*A. psidii*), and Phakopsoraceae (*P. pachyrhizi*). Notably, the three *P. pachyrhizi* isolates—UFV02, K8108, and MT2006—formed a well-supported subclade closely related to *A. psidii* and *Puccinia* species. The second major clade comprised five species, including four representatives of the Melampsoraceae family (*M. allii-populina*, *M. americana*, *M. larici-populina*, and *M. medusae*) and one species from the Coleosporiaceae family (*C. quercuum*).

**Figure 1.**
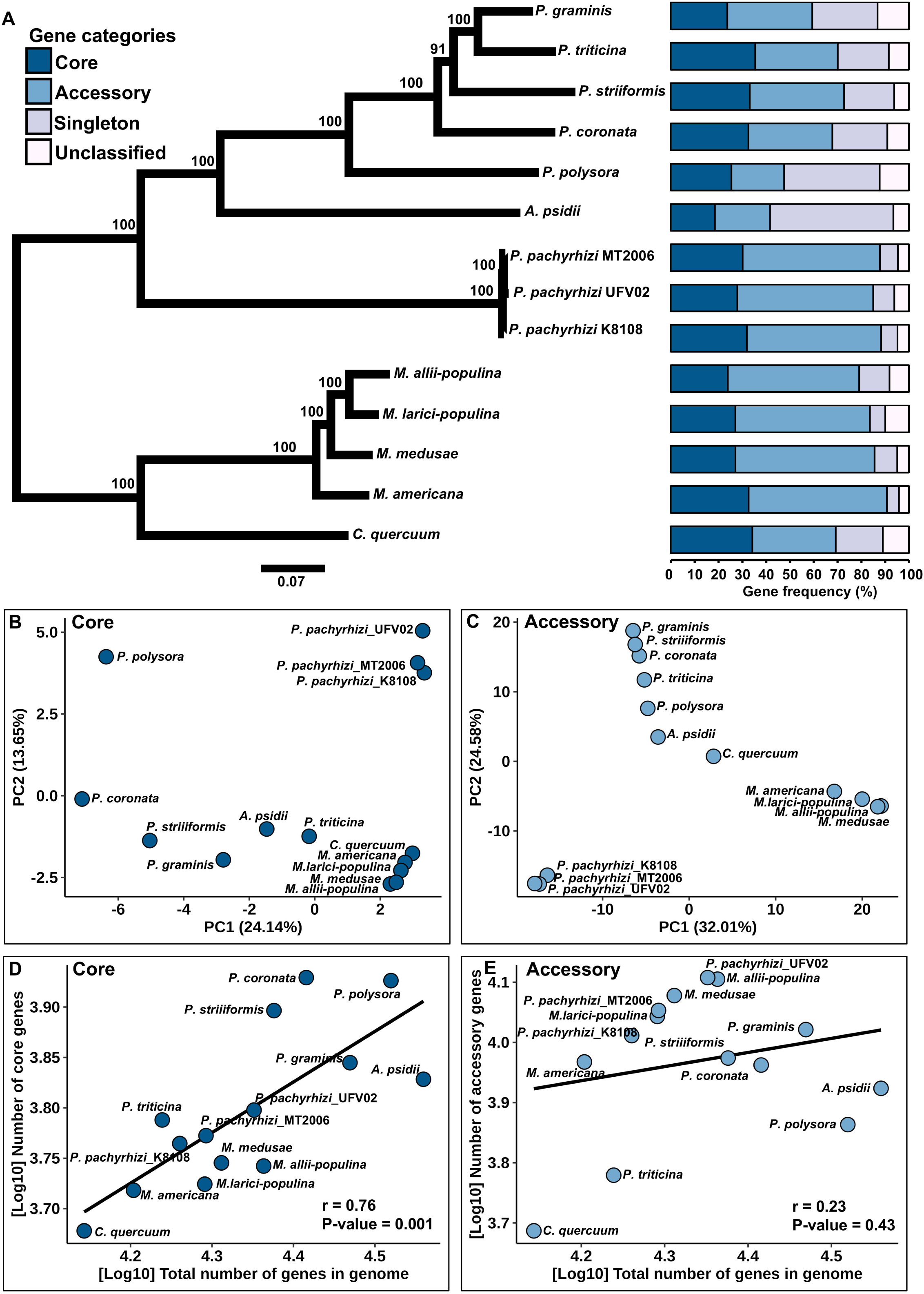
Phylogenomic reconstruction and diversity of core, accessory, and singleton genes across 12 species of the order Pucciniales. **A.** Unrooted maximum likelihood phylogenomic tree based on 44,730 amino acid sites derived from the concatenation of 98 single-copy orthologous proteins. Branch lengths are proportional to the scale bar corresponding to the expected number of substitutions per site. Bootstrap support values are displayed above the branches. Horizontal bar chart showing the relative frequency of core, accessory, and singleton genes per genome. Genes designated as unclassified represent those that were not assigned to any orthogroups according to the OrthoFinder analysis. **B-C.** Principal component analyses (2D PCA) based on the number of core and accessory genes in each orthogroup per genome. **D-E.** Pearson’s linear correlations between [Log_10_] total number of genes per genome and two metrics: [Log_10_] number of core genes; [Log_10_] number of accessory genes. Correlation coefficients (r) and significance levels (P-values) are indicated.

### The size of core, accessory, and singleton gene repertoires in Pucciniales

Our approach, based on OrthoFinder software, identified a highly diverse repertoire of core, accessory, and singleton genes across 12 Pucciniales species. The relative frequency of core genes, calculated as a proportion of core genes within the total number of protein-coding genes, mostly varied from 23.74% (*P. graminis*) to 35.41% (*Puccinia triticina*) (Figure 1A and Supplementary Table S2). One exception was *A. psidii*, which displayed the lowest frequency of core genes at 18.57%. The *P. pachyrhizi* UFV02 reference genome had a frequency of core genes around 28%.

Interestingly, Pucciniales species showed a general tendency to possess larger repertoires of accessory genes than core genes. On average, the relative frequency of accessory genes accounted for 44.61% in each genome. The highest proportions of accessory genes (>55%) were observed in *Melampsora* species and *P. pachyrhizi*. In contrast, lower proportions (<40%) were reported in *A. psidii*, *C. quercuum*, and five *Puccinia* species (Figure 1A and Supplementary Table S2).

The relative frequency of singleton genes varied substantially among the Pucciniales species analyzed (Figure 1A and Supplementary Table S2). Specifically, *Melampsora* species and *P. pachyrhizi* exhibited the lowest frequencies, ranging from 5.07% to 12.75%. Within the Pucciniaceae family, frequencies of singleton genes varied from approximately 21% in *P. striiformis* to about 40% in *P. polysora*. The myrtle rust, *A. psidii*, exhibited the richest singleton gene repertoire, with approximately 52% of its total gene content classified as singleton. For the Pucciniales species, we found no clear association between host species and the size of core, accessory, or singleton repertoires.

### Differences in core and accessory gene composition across the order Pucciniales

To further explore the core gene compositions of the Pucciniales species, we performed a PCA based on the number of core genes in each orthogroup (Figure 1B). The PCA clearly clustered *Melampsora* species and *C. quercuum* together, indicating they shared similar core gene compositions. The PCA-based clustering also revealed that the three *P. pachyrhizi* isolates (UFV02, K8108, and MT2006) formed a single cluster, suggesting they possessed comparable sets of core genes. On the contrary, *Puccinia* species exhibited more varied core gene profiles, with *P. polysora* showing a highly distinct core repertoire (Figure 1B).

Additionally, we conducted a second PCA to examine patterns in accessory gene composition within Pucciniales (Figure 1C), using the number of accessory genes assigned to each predicted orthogroup. This analysis revealed three main clusters, which were congruent with the phylogenomic analysis shown in Figure 1A. One of these clusters involved *Melampsora* species. Meanwhile, the two remaining clusters corresponded to *P. pachyrhizi* and *Puccinia* species, respectively. This clear separation suggested the existence of distinct accessory gene repertoires among *Melampsora*, *P. pachyrhizi* and *Puccinia*, reflecting their distinct evolutionary trajectories within the order Pucciniales.

The estimation of Pearson’s correlation (r) coefficients was calculated to assess relationships between the total number of protein-coding genes and the number of core genes (Figure 1D) or accessory genes (Figure 1E) in Pucciniales genomes. There was a significant correlation between the number of core genes and the total number of protein-coding genes (r = 0.76, P-value = 0.0010; Figure 1D). However, a weak and statistically insignificant correlation was observed for accessory genes (Figure 1E).

### Functional annotation of core, accessory, and singleton genes in *P. pachyrhizi*

Utilizing publicly available functional annotations for three *P. pachyrhizi* isolates, UFV02, K8108, and MT2006 (Gupta et al., 2023), we examined potential biological functions of their core, accessory, and singleton genes. The core and accessory gene pools of *P. pachyrhizi* comprised six major functional categories: CAZymes, proteases, transcription factors, transporters, classically secreted proteins, and putative effector genes (Figure 2). Core and accessory genes associated with secondary metabolism were identified in low numbers, with up to 3-1, respectively. Notably, the secreted proteins, effectors, and transporters were the most well-represented categories in both the core and accessory gene sets (Figure 2). Overall, *P. pachyrhizi* isolates exhibited differences between their core and accessory gene sets, particularly in the composition of secreted proteins, effectors, and transporters. For example, analysis of the *P. pachyrhizi* UFV02 reference genome revealed a considerably higher number of accessory genes than core, specifically for classically secreted proteins (1,235 accessory vs. 542 core genes) and effectors (732 accessory vs. 199 core genes). In contrast, transporters were overrepresented among core genes (537 core vs. 377 accessory genes). The singleton genes of *P. pachyrhizi* were also present among secreted proteins and effectors, although at comparatively lower frequencies (Figure 2).

**Figure 2.**
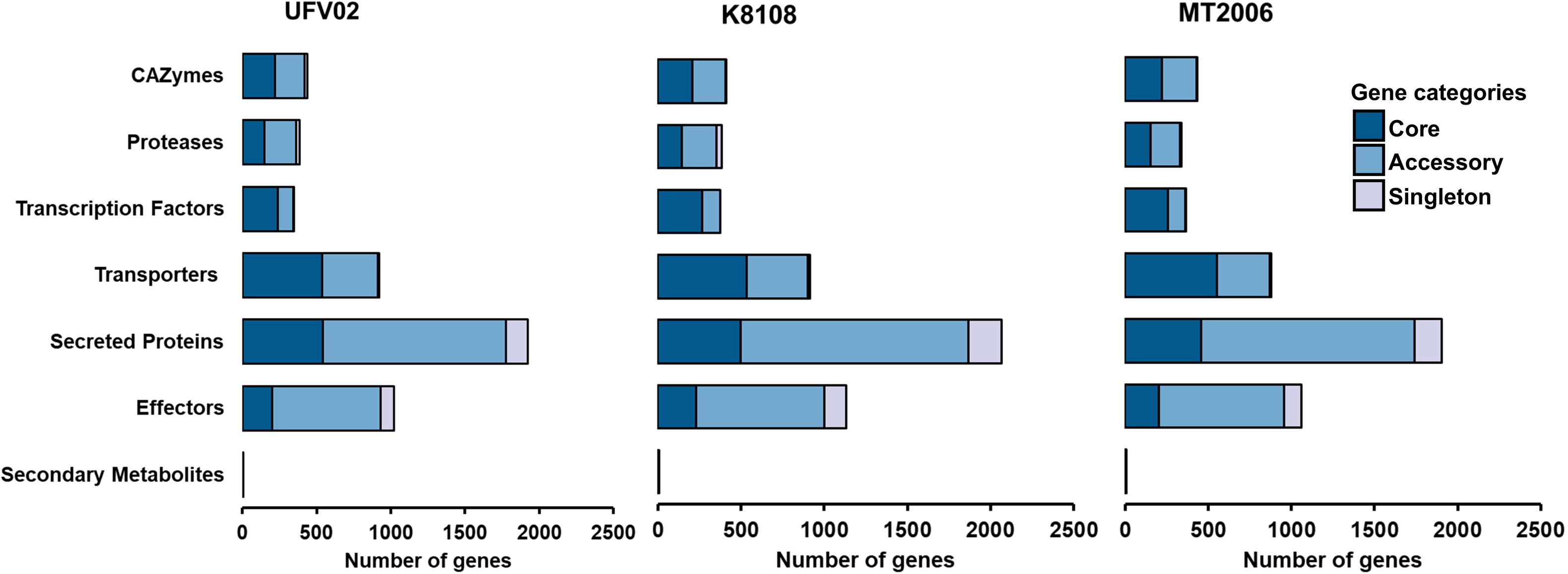
Functional annotation of core, accessory, and singleton genes in three *Phakopsora pachyrhizi* isolates (UFV02, K8108, and MT2006). The vertical axis represents different functional classes: CAZymes, proteases, transcription factors, transporters, classically secreted proteins, effectors, and secondary metabolites. The horizontal bar indicates the number of genes in each functional class.

We performed an in-depth analysis of the gene family compositions within the core, accessory, and singleton compartments of *P. pachyrhizi* (Supplementary Tables S3–S8). Together, core and accessory genes encompassed a total of 79 different CAZyme families, whereas singleton genes were associated with only four families (Supplementary Table S3). The most abundant core CAZyme family was GH5, with up to 21 members per isolate. On the other hand, the AA3 CAZyme family was the largest within the accessory repertoire, with up to 62 members. In total, core and accessory genes displayed 119 PFAM domain annotations related to proteases (Supplementary Table S4). The most notable protease family in the accessory repertoire was Gamma-glutamyl transpeptidase (PF01019), exhibiting up to 66 members. Among transcription factors, 34 predicted PFAM domains were classified as both core and accessory, with a large amount of Zinc finger C2H2 type (PF00096), HMG box (PF00505), and SET domain (PF00856) (Supplementary Table S5). For transporters, the most abundant core family was Sodium/calcium exchanger (PF01699), with a maximum of 22 genes, while the accessory E1–E2 ATPase domain (PF00122) was widely represented, with up to 26 copies (Supplementary Table S6). Numerous transporter families, comprising Mitochondrial carrier proteins (PF00153), Major facilitator superfamily (PF07690), and OPT oligopeptide transporter proteins (PF03169), were identified across both core and accessory compartments. The Amidohydrolase family (PF01979) was the most abundant among accessory secreted proteins (Supplementary Table S7) and accessory effectors (Supplementary Table S8). Other predicted effectors—such as Cellulases (GH5, PF00150), CFEM-domain proteins (PF05730), and GH7 family members (PF00840)—were found in both the core and accessory compartments.

### KOG and GO classification of core, accessory, and singleton genes in *P. pachyrhizi*

The KOG functional classification revealed that the core and accessory genes of *P. pachyrhizi* shared nearly all 25 KOG categories (A-Z) (Figure 3). However, we observed notable differences in the composition of core and accessory genes within KOG categories. For example, the most abundant category J (“Translation, ribosomal structure, and biogenesis”) contained more accessory genes than core genes. In contrast, the opposite pattern was observed in category O (“Post-Translational modification, protein turnover, and chaperones”), in which core genes were predominant. Overall, singleton genes occurred at a low frequency and were primarily found in KOG categories E (Amino acid transport and metabolism) and J.

**Figure 3.**
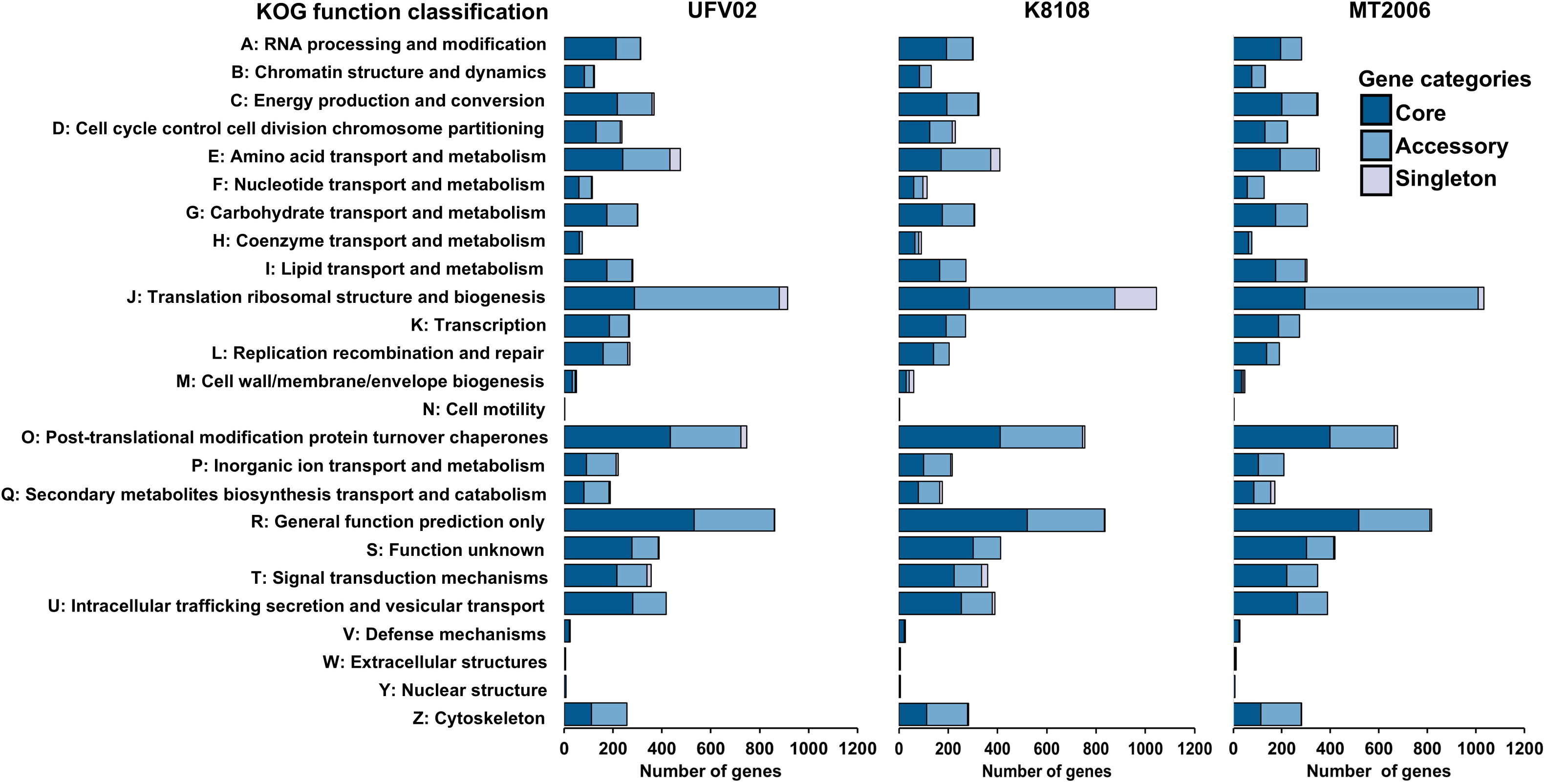
Distribution of Eukaryotic Orthologous Groups (KOG) functional categories among core, accessory, and singleton genes in three *Phakopsora pachyrhizi* isolates (UFV02, K8108, and MT2006). KOG category names are indicated along the vertical axis. The horizontal bar represents the number of genes in each KOG category.

A functional enrichment analysis of KOG categories uncovered that *P. pachyrhizi* accessory genes were significantly enriched in category J, which includes functions related to translation, ribosomal structure, and biogenesis (Supplementary Table S9). Conversely, no KOG categories were significantly enriched among core or singleton genes.

Finally, our GO enrichment analysis identified that certain GO terms were significantly overrepresented in each gene set of *P. pachyrhizi* (Supplementary Table S10). Core genes showed the highest number of enriched GO terms, with 628, followed by accessory genes with 243, and singleton genes with 149 significantly enriched GO terms. In the UFV02 reference genome, the most enriched terms were GO:0009987 (cellular process) for core genes; GO:0005488 (binding) for accessory genes, and GO:0008152 (metabolic process) for singleton genes.

### An expression landscape of core and accessory genes from *P. pachyrhizi*

Based on previous transcriptome resources (Gupta et al., 2023), we characterized the expression levels of core and common accessory genes shared between *P. pachyrhizi* isolates UFV02 and K8108 during soybean rust infection (Figure 4; Supplementary Figure S1). We focused only on the top differentially expressed genes (DEGs) with a Log2 fold change of at least 3 (see the Material and Methods section). The gene expression analysis was performed across distinct functional groups, representing putative effectors, CAZymes, proteases, transcription factors, and transporters.

**Figure 4.**
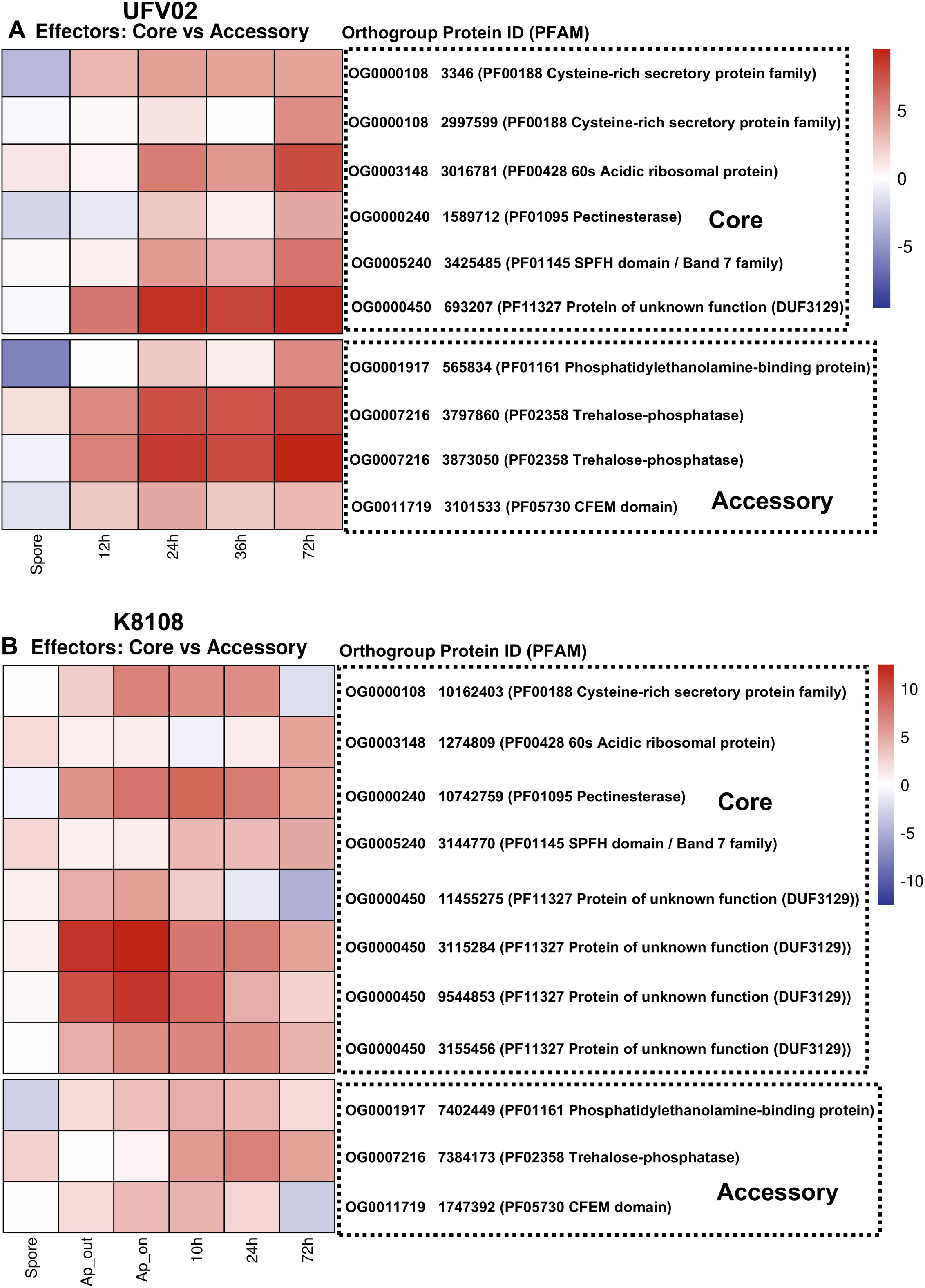
Heatmap depicting expression profiles of common core and accessory effector top DEGs shared between *Phakopsora pachyrhizi* isolates (UFV02 and K8108) during soybean infection. Expression levels were evaluated under both *in-vitro* and *in-planta* conditions. The *in-vitro* stages included spore and appressorium samples (Ap_out), while *in-plant* stages spanned 10 to 72 hours post-inoculation. The *in-plant* appressorium samples (Ap_on) were exclusively evaluated for isolate K81081. Expression levels were calculated as Log2 fold change (Log2FC). The x-axis represents distinct time points at which gene expression was evaluated. The y-axis represents the common core and accessory effector top DEGs shared between UFV02 and K8108 isolates. Color intensity reflects expression levels according to the color palette. Gene expression data were derived from RNA-seq libraries analyzed in a previous study (Gupta et al., 2023).

Notably, both core and common accessory effectors were intensely upregulated during appressorium formation and infection stages (Figure 4). One interesting finding was that different PFAM domains were observed between expressed core and accessory effectors. Overall, expressed core effectors contained five PFAM domains: PF00188 (cysteine-rich secretory protein), PF00428 (60S acidic ribosomal protein), PF01095 (pectinesterase), PF01145 (SPFH domain/Band 7), and PF11327 (protein of unknown function, DUF3129). In contrast, common accessory effectors harbored distinct PFAM domains, such as PF01161 (phosphatidylethanolamine-binding protein), PF02358 (trehalose-phosphatase), and PF05730 (CFEM domain).

Among the core effectors, expression levels varied substantially, with genes containing the PFAM domain PF11327 exhibiting the highest expression abundance, while the remaining genes displayed comparatively lower expression (Figure 4). Within the set of common accessory effectors, genes harboring the PFAM domain PF02358 were strongly induced, particularly in the UFV02 isolate (Figure 4A).

Expression analyses of CAZymes revealed that core genes from several families (e.g., CE4, EXPN/CBM63, GH5, GH7, and GH131) were highly expressed during soybean infection (Supplementary Figure 1A–B). Within the GH18 chitinase family, both core and common accessory genes were strongly upregulated. Among common accessory families, GH79, GH26, and GT10 also displayed elevated expression levels.

Proteases were considerably upregulated during infection, with the UFV02 isolate reaching a maximum of 33 expressed proteases across core and accessory repertoires (Supplementary Figure 1C–D). The highest expression levels were observed in two core protease families (PF00026: eukaryotic aspartyl protease; PF00450: serine carboxypeptidase) and one common accessory family (PF12697: alpha/beta hydrolase).

The transporter category contained the highest number of expressed core and accessory genes, with a total of 120 in the UFV02 isolate (Supplementary Figure 1E–F). The most upregulated core transporters were associated with various PFAM domains, including PF00230 (major intrinsic protein), PF00324 (amino acid permease), PF01794 (ferric reductase–like transmembrane protein), PF02096 (60 kDa inner membrane protein), and PF09084 (NMT1/THI5 proteins). Both OPT oligopeptide transporters (PF03169) and major facilitator superfamily genes (PF07690) exhibited robust expression within the core and common accessory sets.

A low number of core transcription factors was induced during the soybean disease cycle, with a maximum of 11 in the UFV02 isolate (Supplementary Figure 1G–H). The most highly expressed core transcription factors were annotated as bZIP (PF00170), GATA zinc finger (PF00320), STE-like (PF02200), and homeobox KN (PF05920). No highly expressed accessory transcription factor was identified under our selection criteria.

### Molecular diversity of core and accessory effectors from *P. pachyrhizi*

We examined genetic variation among orthogroups containing core and common accessory effectors previously included in our expression analyses (Figure 4). Distinct levels of genetic diversity were observed among effector orthogroups from *P. pachyrhizi* (Table 1). Specifically in the core effector orthogroups, OG0000108 (representing cysteine-rich secretory protein family) exhibited the highest diversity (nucleotide diversity, π ≈ 0.34; Watterson’s Theta, θw ≈ 0.27), while OG0003148 (60s acidic ribosomal protein) and OG0005240 (SPFH domain/Band 7 family) showed no detectable polymorphism among their members. In cases of accessory effectors, we identified elevated genetic differentiation in orthogroups OG0001917 (Phosphatidylethanolamine-binding proteins) and OG0007216 (Trehalose-phosphatase), with π ≈ 0.31 and π ≈ 0.25, respectively.

**Tabela 1.**
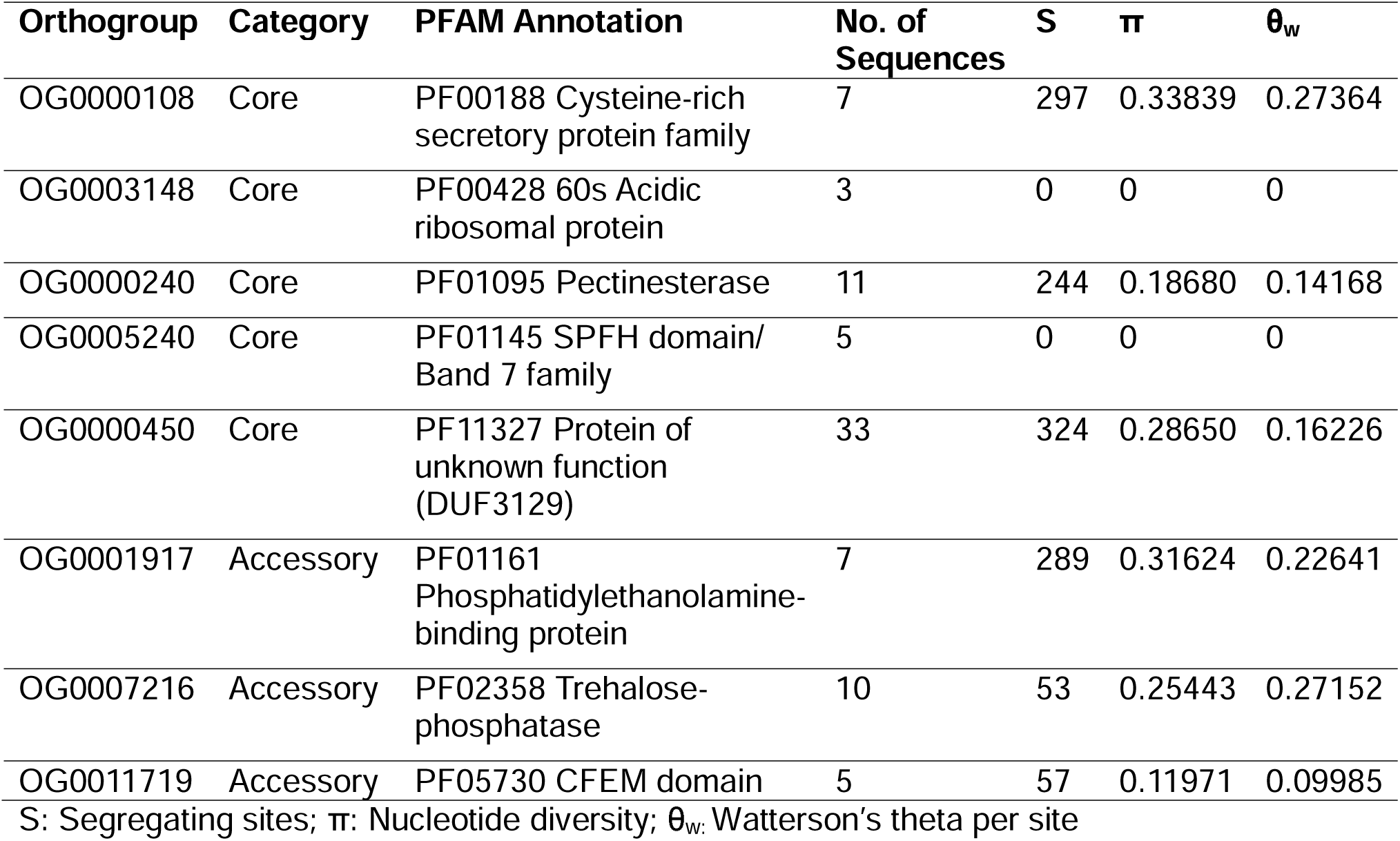
Measures of molecular diversity for core and accessory orthogroups in *Phakopsora pachyrhizi.* Each orthogroup included at least one top differentially expressed gene (DEG), which was identified as either a core effector or a common accessory effector shared by both UFV02 and K8108 isolates.

## DISCUSSION

Our results uncovered that the accessory gene repertoire of Pucciniales species is relatively large. On average, accessory genes accounted for 44.61% of the total gene content in Pucciniales, with some species such as *P. pachyrhizi* reaching approximately 57% (Supplementary Table S2). This proportion of accessory genes in Pucciniales is substantially higher than that reported for other widely distributed fungal plant pathogens with biotrophic or hemibiotrophic lifestyles. For instance, the hemibiotrophic wheat pathogen *Zymoseptoria tritici* contains approximately 30% of accessory gene content (Badet et al., 2020), while the biotrophic ergot fungus *Claviceps purpurea* harbors about 27.05% (Wyka et al., 2022).

In addition to their large repertoires of accessory genes, Pucciniales species also displayed notable variation in the distribution of accessory genes. Our findings showed that *Melampsora* species and *P. pachyrhizi* have the highest accessory content, whereas *A. psidii*, *C. quercuum*, and several *Puccinia* species possess comparatively smaller accessory components (Supplementary Table S2). These differences likely reflect distinct evolutionary trajectories across Pucciniales lineages. Gene duplication events may have contributed to the expansion of accessory repertoires in *P. pachyrhizi* and *Melampsora* species. On the other hand, gene loss may help explain the reduced accessory gene content in *A. psidii*, *C. quercuum*, and *Puccinia* species. In fact, previous evidence has documented lineage-specific patterns of gene gain and loss shaping the genomes of multiple rust fungi, including *C. quercuum*, *M. larici-populina*, *P. graminis* (Pendleton et al., 2014), and *P. pachyrhizi* (Gupta et al., 2023).

As expected, the compositions of accessory orthogroups (Figure 1C) closely mirrored the phylogenomic placements among Pucciniales species (Figure 1A). The observed concordance suggests that the diversification of accessory gene repertoires occurred independently within each Pucciniales species, likely in response to distinct host-related selective pressures.

In general, Pucciniales genomes with higher numbers of genes also show larger core repertoires, as shown by a significant positive correlation between core gene content and the total number of protein-coding genes (Figure 1D). This pattern suggests that core components were largely retained after gene expansion events within Pucciniales genomes. Given their crucial roles in cellular metabolism, reproduction, and other essential biological processes (Croll & McDonald, 2012), the retention of core genes is expected.

In contrast to core genes, the size of the accessory repertoire showed no significant correlation with the total number of genes in Pucciniales (Figure 1E). The absence of correlation indicates that accessory genes of Pucciniales have experienced extensive evolutionary dynamics, characterized by frequent gains and losses of gene families. Consistent with the TE-rich genomes of Pucciniales (Corre et al., 2025), the evolution of accessory gene content in Pucciniales may be influenced by driving factors such as gene duplications or deletions associated with TE activity. Some species across Pucciniales exhibit the highest levels of TE content in Basidiomycota, for example, in *Melampsora* species, TE content can achieve more 50% of the genome, while in *P. pachyrhizi* it can cover more than 90% of the genome (Corre et al., 2025). The impact of TEs on accessory genes has been examined in fungal pathogens. For instance, accessory genes in the maize pathogen *Cercospora zeina* were found in proximity to TEs (Welgemoed et al., 2023). In the plant-pathogenic fungus *Claviceps purpurea*, TE-mediated duplications have contributed to the expansion of accessory orthogroups (Wyka et al., 2022). Another evolutionary fate of many accessory genes in Pucciniales is eventual loss or pseudogenization due to their tendency to accumulate rapid sequence changes, including loss-of-function mutations, premature stop codons, and deleterious frameshifts, as observed in *Zymoseptoria tritici* (Plissonneau et al., 2018)

Through bioinformatic approaches, our study provides the first insight into the potential functional relevance of core and accessory effector genes in *P. pachyrhizi*. Consistent with a previously reported high abundance of effectors in *P. pachyrhizi* genomes (Gupta et al., 2023), this functional category was one of the most well-represented in both the core and accessory gene sets (Figure 2). The widespread presence of effectors across the core and accessory genomic compartments of *P. pachyrhizi* highlights their adaptive significance to the pathogen’s ability to subvert host defense responses.

Our expression analyses revealed strong upregulation of both core and accessory effector genes during the early and late stages of *P. pachyrhizi* infection on soybean (Figure 4). Several of these highly expressed effectors contained PFAM domains previously associated with fungal pathogenicity. Among them, the PF11327 domain was identified in significantly induced core effectors. This domain has been reported in rust fungal effectors (Saunders et al., 2012; Castanho et al., 2024). In *P. pachyrhizi*, PF11327 occurs in Egh16-like effectors capable of repressing pathogen-associated molecular pattern–triggered immunity (PTI) mechanisms, including the suppression of reactive oxygen species (ROS) production and callose deposition (Castanho et al., 2024). The cysteine-rich PF00188 domain was also detected in core effectors and showed elevated expression, especially in the isolate UFV02 (Figure 4A). While the functional role of PF00188 remains uncharacterized in rust fungi, it has been linked to virulence in the ascomycete fungus *Fusarium oxysporum* (Prados-Rosales et al., 2012). Among the expressed core effectors of *P. pachyrhizi*, another notable pathogenicity-related domain was pectinesterase (PF01095). The secretion of core effector pectinesterases in *P. pachyrhizi* likely facilitates fungal infection by mediating pectin degradation, a major component of the plant cell wall. The *in-planta* upregulation of pectinesterase genes has also been reported in phytopathogenic organisms, including *Corynespora cassiicola* (Rocha et al., 2023) and *Phytophthora capsici* (Li et al., 2011). We also found that the core effector SPFH/Band 7 genes of *P. pachyrhizi* exhibited increased expression, a finding consistent with their multifaceted biochemical roles. Indeed, a body of evidence suggests that SPFH/Band 7 proteins mediate a diverse array of molecular functions, including hyphal growth, mitochondrial activity, and stress-response signaling in pathogenic fungi (Heredia & Rauceo, 2021).

Notably, phosphatidylethanolamine-binding protein family members, classified as accessory effectors, showed elevated transcriptional profiling in *P. pachyrhizi*. Phosphatidylethanolamine-binding proteins are typically abundant in rust fungal secretomes and appear to be involved in lipid binding, serine protease inhibition, and regulation of the MAP kinase pathway (Saunders et al., 2012). In addition, we documented that accessory effector trehalose-phosphatases (PF02358) of *P. pachyrhizi* also exhibited robust expression, suggesting a potential role in pathogenesis. This observation is consistent with virus-induced gene silencing (VIGS) assays in the bean rust fungus *Uromyces appendiculatus*, which demonstrated that a trehalose-phosphatase candidate effector contributes significantly to pathogenicity (Cooper & Campbell, 2017). The induced expression of accessory effectors containing a Common Fungal Extracellular Membrane (CFEM) domain in *P. pachyrhizi* UFV02 is coherent with previous functional data of wheat stripe rust (*Puccinia striiformis* f. sp. *tritici*; *Pst*), where the putative effector PstCFEM1 enhances virulence by suppressing host cell death, callose deposition, and the accumulation of ROS (Bai et al., 2022).

The identification of the 60s acidic ribosomal protein domain (PF00428) among the upregulated core effectors of *P. pachyrhizi* is an intriguing finding. It remains unclear how core effectors encoding a predicted 60s acidic ribosomal protein domain may contribute to the infection process, despite their marked upregulation during late stages of the infection time course (Figure 4). The lack of functional data for these genes precluded further inferences regarding their potential biological role.

Only a few functional studies have been experimentally validated the biological importance of *P. pachyrhizi* effectors during host–pathogen interactions (De Carvalho et al., 2017; Qi et al., 2018; Bueno et al., 2022). These investigations showed that *P. pachyrhizi* candidate effectors can suppress effector-triggered immunity (ETI) by inhibiting the hypersensitive response (De Carvalho et al., 2017) as well as induce PTI suppression (Bueno et al., 2022; Castanho et al., 2024). In our analysis, we discovered that several previously characterized *P. pachyrhizi* effectors corresponded to highly conserved core genes across rust fungi (data not shown). For example, two validated effectors (de_novo_1784 and de_novo_2238) described by De Carvalho et al., (2017) and Castanho et al., (2024) corresponded to expressed core genes clustered within orthogroup OG0000450 with an Egh16-like domain. The functionally validated effector Phapa-7431740, which harbors a calcium-binding EGF domain (Bueno et al., 2022), was assigned to the core orthogroup OG0000034. Several *P. pachyrhizi* effectors analyzed by Qi et al., (2018) displayed high sequence similarity to genes allocated into multiple core and accessory orthogroups in our dataset. Additionally, computational prediction analyses indicated that a putative accessory effector phosphatidylethanolamine-binding protein (UFV02-3006262) from *P. pachyrhizi*, which belongs to the orthogroup OG001917, may interact with an R-protein localized at the soybean rust resistance *Rpp1* locus (Sodrzeieski, unpublished data).

Although *P. pachyrhizi* displays low intra-lineage genetic diversity with predominantly clonal reproduction (Rocha et al., 2025), its virulence-associated loci likely undergo rapid diversification to facilitate host adaptation. Since effector gene families in plant-pathogens are typically characterized by high levels of genetic variation (Dal’Sasso et al., 2022; Rocha et al., 2023), it is plausible that specific effector loci in *P. pachyrhizi* may experience accelerated evolutionary rates to overcome host resistance despite a clonal population structure. Our findings demonstrate that core and accessory effector orthogroups of *P. pachyrhizi* exhibit distinct degrees of polymorphism, suggesting they are subject to divergent evolutionary pressures (Table 1). The absence of genetic polymorphism in core effector orthogroups OG0003148 (60s acidic ribosomal protein) and OG0005240 (SPFH domain/Band 7 family) likely reflects strong purifying selection. By eliminating deleterious mutations, purifying selection preserves genes with essential functions, such as core effectors, thereby reducing allelic variation across *P. pachyrhizi* populations. Conversely, genomic rearrangements and sequence modifications, coupled with relaxed selective pressure, may drive the high diversification observed in the cysteine-rich core effector orthogroup OG0000108 and accessory effector orthogroups OG0001917 (Phosphatidylethanolamine-binding proteins) and OG0007216 (Trehalose-phosphatase). Given their high genetic variability and robust expression during soybean infection, members of these effector orthogroups might have experienced quick genetic diversification, generating alternative alleles to allow rapid adaptive changes throughout the cycle of soybean rust disease.

Our findings support that multiple core and accessory gene families of CAZymes, proteases, transcription factors, and transporters are involved in the *P. pachyrhizi*-soybean interactions. The coordinated expression of these genes likely has driven the successful colonization of the host by *P. pachyrhizi*.

CAZymes are commonly secreted by plant pathogenic fungi to degrade host cell wall components, thereby facilitating the penetration of leaf tissues (Zhao et al., 2013). It is believed that *P. pachyrhizi* utilizes degrading enzymes to breach the host cuticle and penetrate the epidermal cell wall, enabling primary hyphal growth and subsequent haustorium development (Edwards & Bonde, 2001). Our results showed that a set of core CAZymes in *P. pachyrhizi* was upregulated during soybean infection, including cell wall–degrading enzymes such as cellulases (GH5 and GH7), cutinases (CE5), and pectinesterases (CE8) (Supplementary Figure 1A–B). Similarly, GH5, GH7, and CE5 families exhibited elevated transcriptional levels in *M. larici-populina* during poplar rust infection (Duplessis et al., 2011). In agreement with previous observations that rust fungi possess an expanded repertoire of GH5 cellulases (Zhao et al., 2013), we found that GH5 represents the most abundant core CAZyme family in *P. pachyrhizi*.

Remarkably, AA3 emerges as the largest accessory CAZyme family in *P. pachyrhizi* (Supplementary Table S3). AA3s belong to the glucose–methanol–choline (GMC) oxidoreductase superfamily and may act as ligninolytic enzymes (Sützl et al., 2019). A significant expansion of AA3 GMC oxidoreductases was reported in *P. pachyrhizi* (Gupta et al., 2023).

As observed in *P. pachyrhizi* transcriptomes, core genes encoding CE4 chitin deacetylases and GH18 chitinases were strongly induced at multiple time points during interaction with the soybean host. One accessory member of the GH18 chitinase family also exhibited high expression during advanced stages of infection. These findings are consistent with previously reported expansions of the GH18 chitinase family in *P. pachyrhizi* (Gupta et al., 2023). Expression of chitinase genes has also been documented in wheat stripe rust, *P. striiformis* f. sp. *tritici*, particularly in haustoria (Garnica et al., 2013). In fungi, chitinases have been proposed to be involved in diverse functions, including fungal cell wall remodeling, hyphal growth, nutrient acquisition strategies, mycoparasitism, and suppression of plant chitin receptors to evade host immune responses (Langner & Göhre, 2015).

Fungal proteases are frequently required for pathogenicity, nutrient acquisition, and the degradation of plant immune receptors (Chandrasekaran et al., 2016). In line with previous transcriptomic analyses of rust fungi (Duplessis et al., 2011; Garnica et al., 2013), our study identified a specific subset of proteases that were markedly upregulated during the biotrophic interaction between *P. pachyrhizi* and soybean. This subset encompasses core proteases (aspartyl proteases and serine carboxypeptidases), as well as accessory proteases (alpha/beta hydrolases) (Supplementary Fig. 1C–D). We hypothesized that these core and accessory proteases might confer adaptive advantages by suppressing host immune mechanisms, thus promoting *P. pachyrhizi* pathogenicity. Indeed, earlier molecular characterization supports the importance of proteases in plant-pathogen interactions. For example, an aspartic protease effector, PpEC15, from *P. pachyrhizi* demonstrates the ability to repress basal plant defenses (Chicowski et al., 2023). This effector interacts specifically with two soybean proteins — the peptide chain release factor *Gm*PCRF and the transcription factor *Gm*NAC83 — and cleaves *Gm*DAHP, a key enzyme in the shikimate pathway (Chicowski et al., 2023). Moreover, fungal serine carboxypeptidases are known to influence growth, morphology, stress tolerance, and virulence (Zhang et al., 2022; Umer et al., 2025),

Due to their biotrophic lifestyle, rust fungi rely on host plants as a source of nutrients in order to obtain carbohydrates and amino acids (Struck, 2015; Lorrain et al., 2019). Nutrient acquisition in rust fungi is believed to involve a suite of transporter proteins, which generally exhibit high and specific expression in haustoria during biotrophic growth (Lorrain et al., 2019). Among rust fungi, only a limited number of transporter proteins have been functionally characterized, including amino acid permeases (AAT1p, AAT2p, and AAT3p), the proton pump H+-ATPase, and the hexose transporter HXT1 (Struck, 2015). Here, we identified both core and accessory transporter genes of *P. pachyrhizi* that were upregulated during soybean rust infection (Supplementary Figure 1E–F). Highly expressed core transporters in *P. pachyrhizi* included genes involved in amino acid transport (e.g., amino acid permeases, PF00324) and iron assimilation, particularly ferric reduction activity (e.g., ferric reductase-like transmembrane proteins, PF01794). The co-expression of core and accessory major facilitator superfamily (MFS) transporters indicates that they might be required for nutrient uptake in *P. pachyrhizi* infection. The MFS family constitutes a ubiquitous group of transmembrane transporters that is responsible for catalyzing the transport of a variety of molecules, such as nucleosides, amino acids, short peptides, and lipids (Yan 2015). Similar expression patterns of MFS transporters have been reported in other rust fungi (Duplessis et al., 2011; Garnica et al., 2013). As expected, we detected that the oligopeptide transporters (OPT) core and accessory genes were overexpressed in *P. pachyrhizi*. A previous study showed that the OPT gene family is significantly expanded in Pucciniales and is expressed in poplar rust *M. larici-populina*, indicating a specialized role in nitrogen and sulfur uptake during host infection (Guerillot et al., 2023).

The upregulation of core transcription factors in *P. pachyrhizi* likely reflects their multifunctional roles in regulating fungal development and metabolism. The core set of differentially expressed transcription factors in *P. pachyrhizi* consisted predominantly of bZIP, GATA zinc finger, STE-like, and homeobox KN families. These transcription factors are widely distributed in fungi and govern complex biological processes associated with fungal growth, conidiation, and host-pathogen interactions (Gu et al., 2014; Calvo et al., 2024; Hu et al 2025). For instance, bZIP transcription factors have been implicated in the regulation of mycelial growth, pathogenicity, sulfur metabolism, iron homeostasis, and responses to oxygen and osmotic stress (Kong et al., 2015; Zhao et al., 2022).

In conclusion, this study offers a comprehensive overview of the core and accessory genomic components in rust fungi. Overall, Pucciniales genomes harbor a large and evolutionarily dynamic accessory gene repertoire, likely shaped by recurrent gene expansions and contractions. In contrast, the core genomic component displays a comparatively small size across most Pucciniales species. Notably, core genes regulate vital biological processes and are therefore prominently retained throughout the evolution of Pucciniales. In *P. pachyrhizi*, numerous core and accessory families—encoding effectors, CAZymes, proteases, transporters, and transcription factors—were strongly expressed, underscoring their contribution to successful soybean rust infection. The highly upregulated core effectors of *P. pachyrhizi* are characterized by cysteine-rich proteins, pectin-degrading enzymes, and SPFH/Band 7 proteins, whereas the accessory effectors encode phosphatidylethanolamine-binding protein, trehalose-phosphatases, and CFEM domain. Our work provides a foundation for prioritizing candidate genes to support future functional studies aimed at uncovering the molecular mechanisms of adaptation in *P. pachyrhizi*.

## AUTHOR CONTRIBUTIONS

**Vinicius Delgado da Rocha**: Conceptualization, Formal analysis, Visualization, Writing – original draft. **Liliane Santana Oliveira:** Identification of redundant proteins and manuscript revision. **Francismar Corrêa Marcelino-Guimarães**: Conceptualization, Funding Acquisition, Supervision, Writing – reviewing and editing. All authors read and approved the submitted version.

## ACKNOWLEDGMENTS

We are grateful to the funding agency, the National Council of Scientific and Technological Development - CNPq (project 440782/2022-8 and DTI-A 385056/2024-9). We thank Dr. Peri Tobias (University of Sidney) for kindly providing proteome data from *A. psidii*.

## AI LANGUAGE MODEL ASSISTANCE

ChatGPT tool was exclusively used for language polishing services under the author’s supervision. We are entirely responsible for the manuscript content.

## DATA AVAILABILITY

All genomic data used in this study were obtained from public sources. Additional details are provided in Supplementary Table S1.

