## Supplementary Figure 1 for "COMPARATIVE GENOMIC ANALYSIS OF CORE AND ACCESSORY GENES IN RUST FUNGI REVEALS PATHOGENICITY-ASSOCIATED GENE FAMILIES IN *Phakopsora pachyrhizi*"

### Supplementary Figure S1

**Supplementary Figure S1**. Heatmap showing top differentially expressed genes (DEGs) classified as core or common accessory shared between two *Phakopsora pachyrhizi* isolates (UFV02 and K8108) during soybean infection. Gene expression was assessed across different functional classes: (A–B) CAZymes, (C–D) proteases, (E–F) transporters, and (G–H) transcription factors. Expression levels were evaluated under both *in-vitro* and *in-planta* conditions. The *in-vitro* stages included spore and appressorium samples (Ap_out), while *in-plant* stages spanned 10 to 72 hours post-inoculation. The *in-plant* appressorium samples (Ap_on) were exclusively evaluated for isolate K81081. Expression levels were calculated as Log2 fold change (Log2FC). The x-axis represents distinct time points at which gene expression was evaluated, while the y-axis represents the common core and accessory top DEGs shared between UFV02 and K8108 isolates. Protein IDs and PFAM domains are indicated along the y-axis. Color intensity reflects expression levels according to the color palette. Gene expression data were derived from RNA-seq libraries analyzed in a previous study (Gupta et al., 2023).

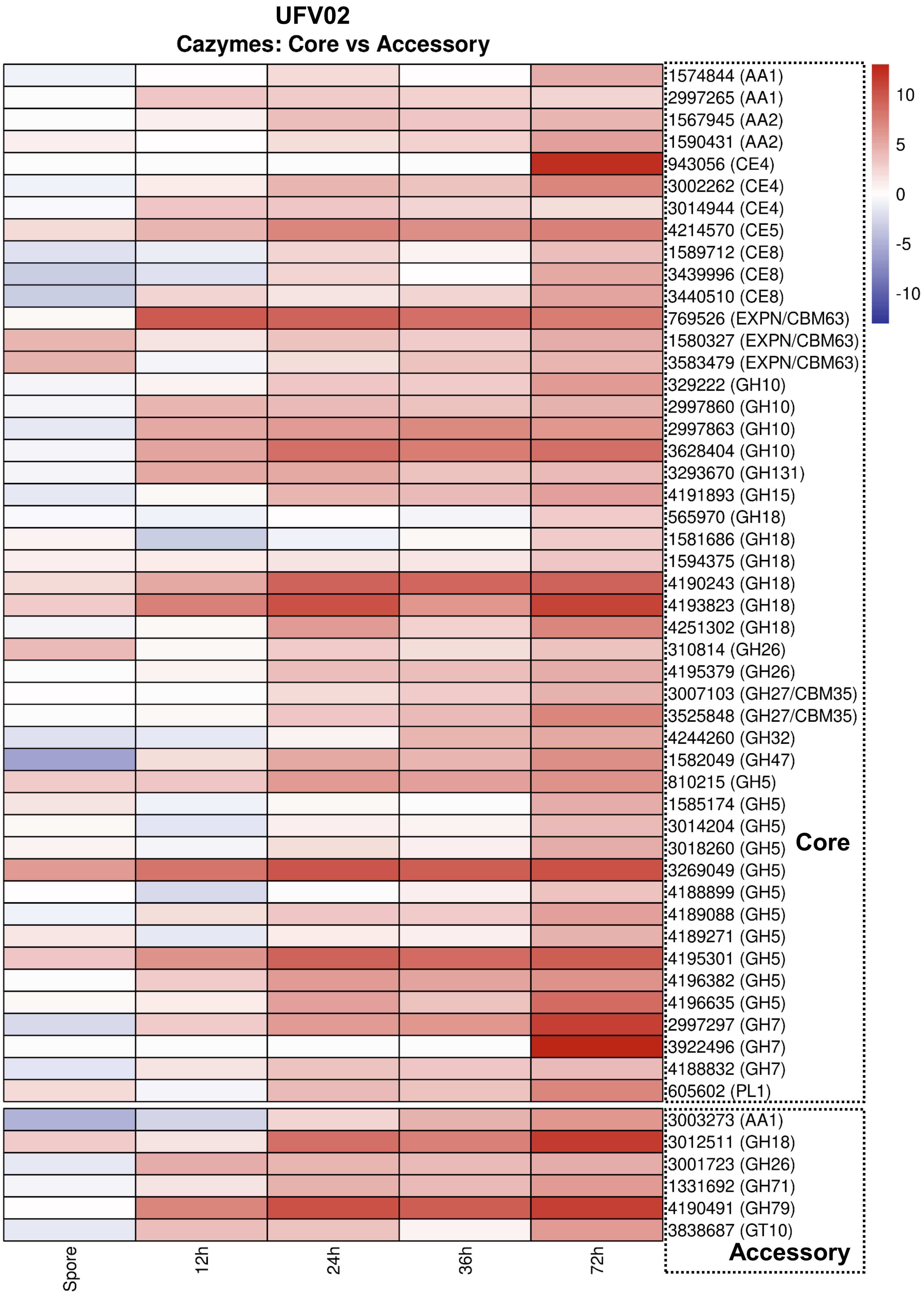


**Supplementary Figure S1-A**

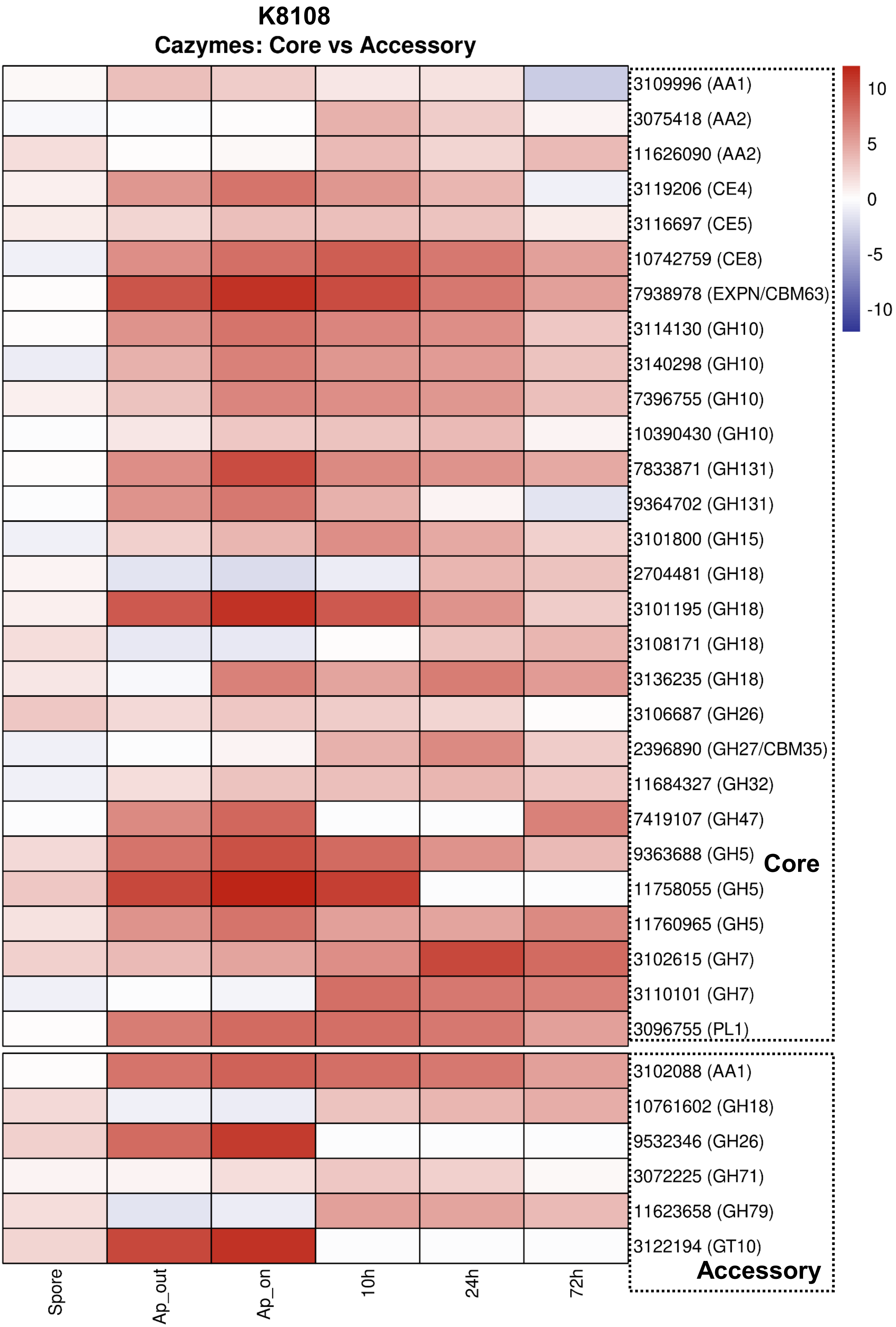


**Supplementary Figure S1-B**

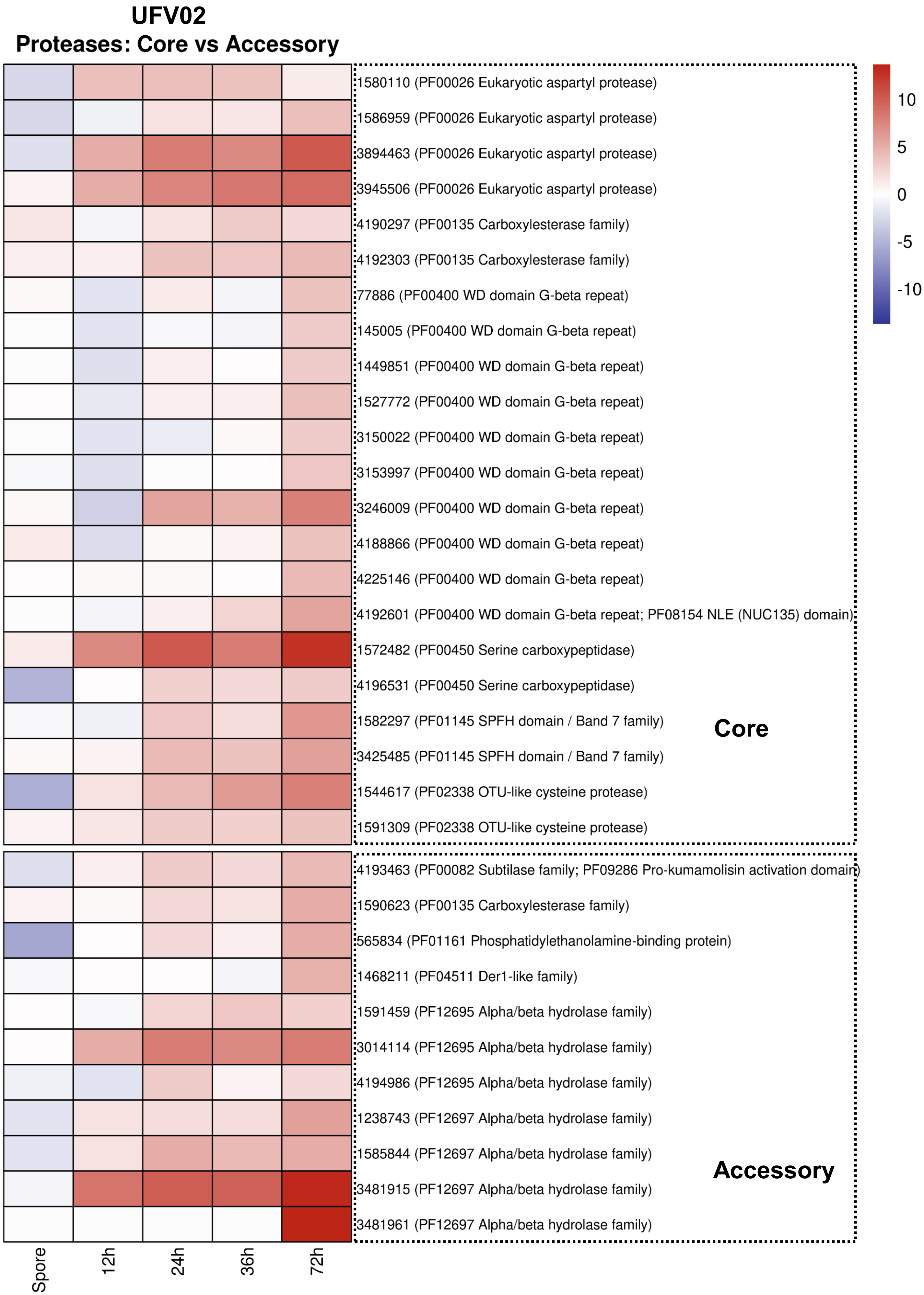


**Supplementary Figure S1-C**

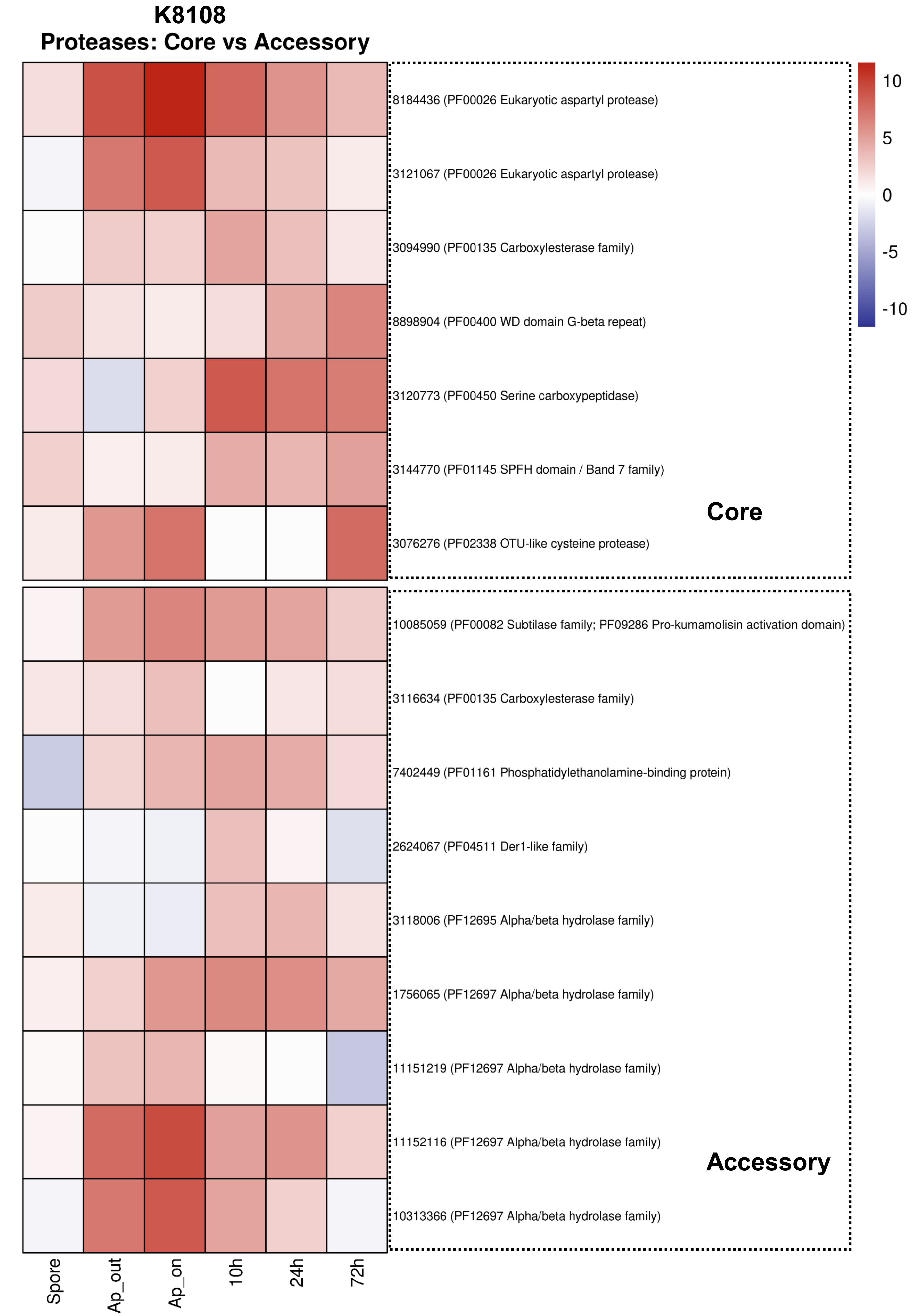


**Supplementary Figure S1-D**

#### UFV02
#### Transporters: Core vs Accessory

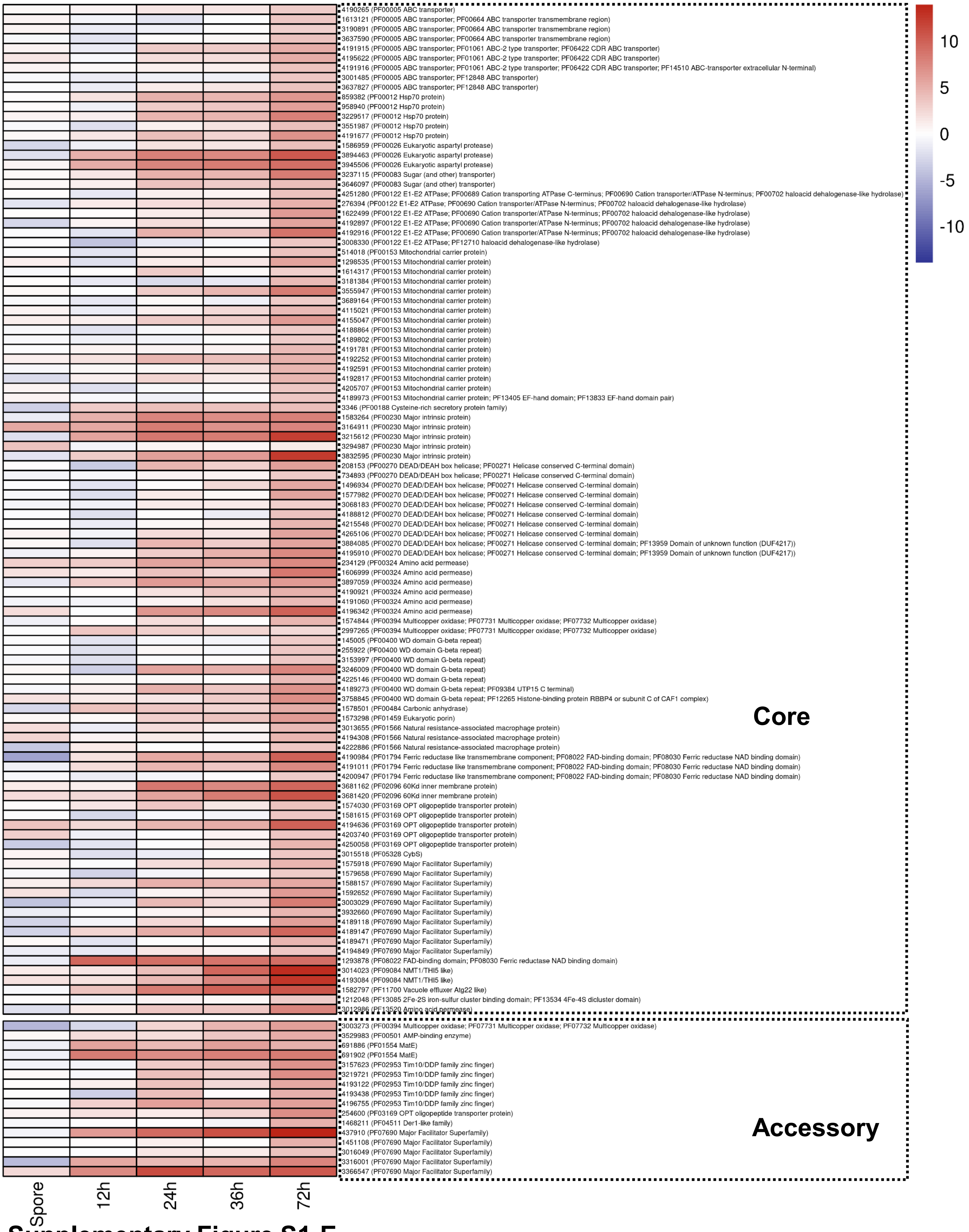


**Supplementary Figure S1-E**

## K8108
#### Transporters: Core vs Accessory

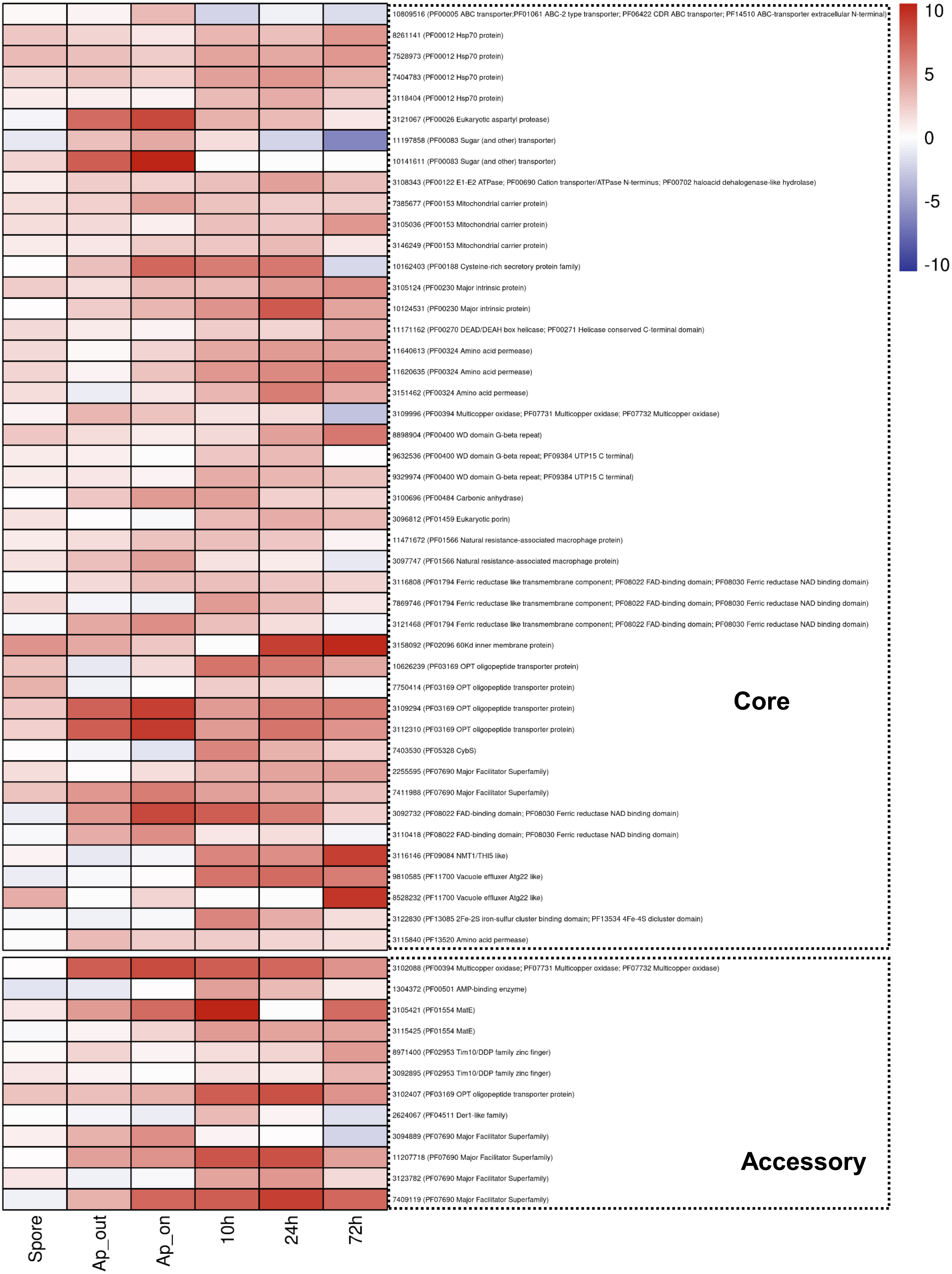


**Supplementary Figure S1-F**

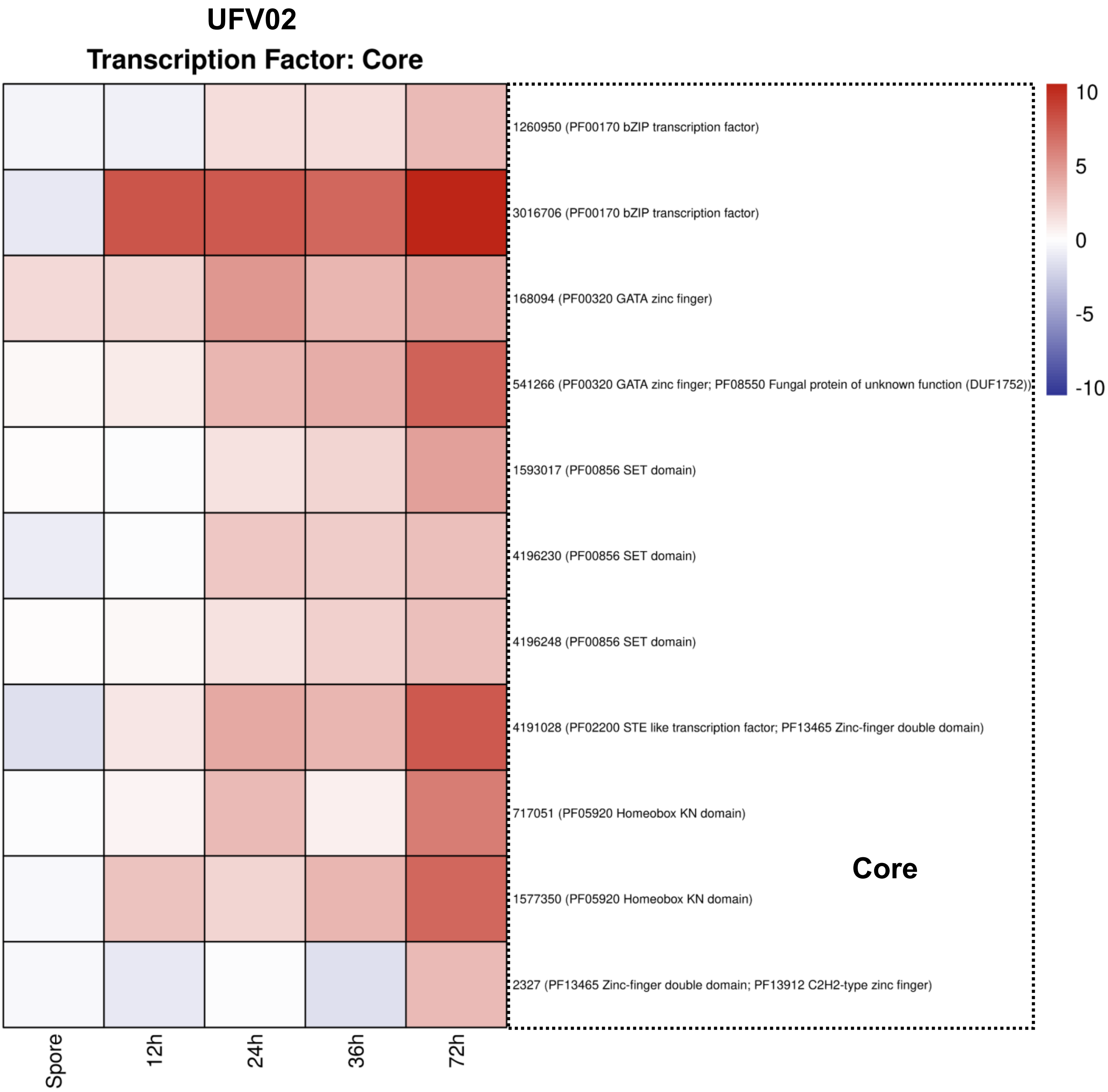


**Supplementary Figure S1-G**

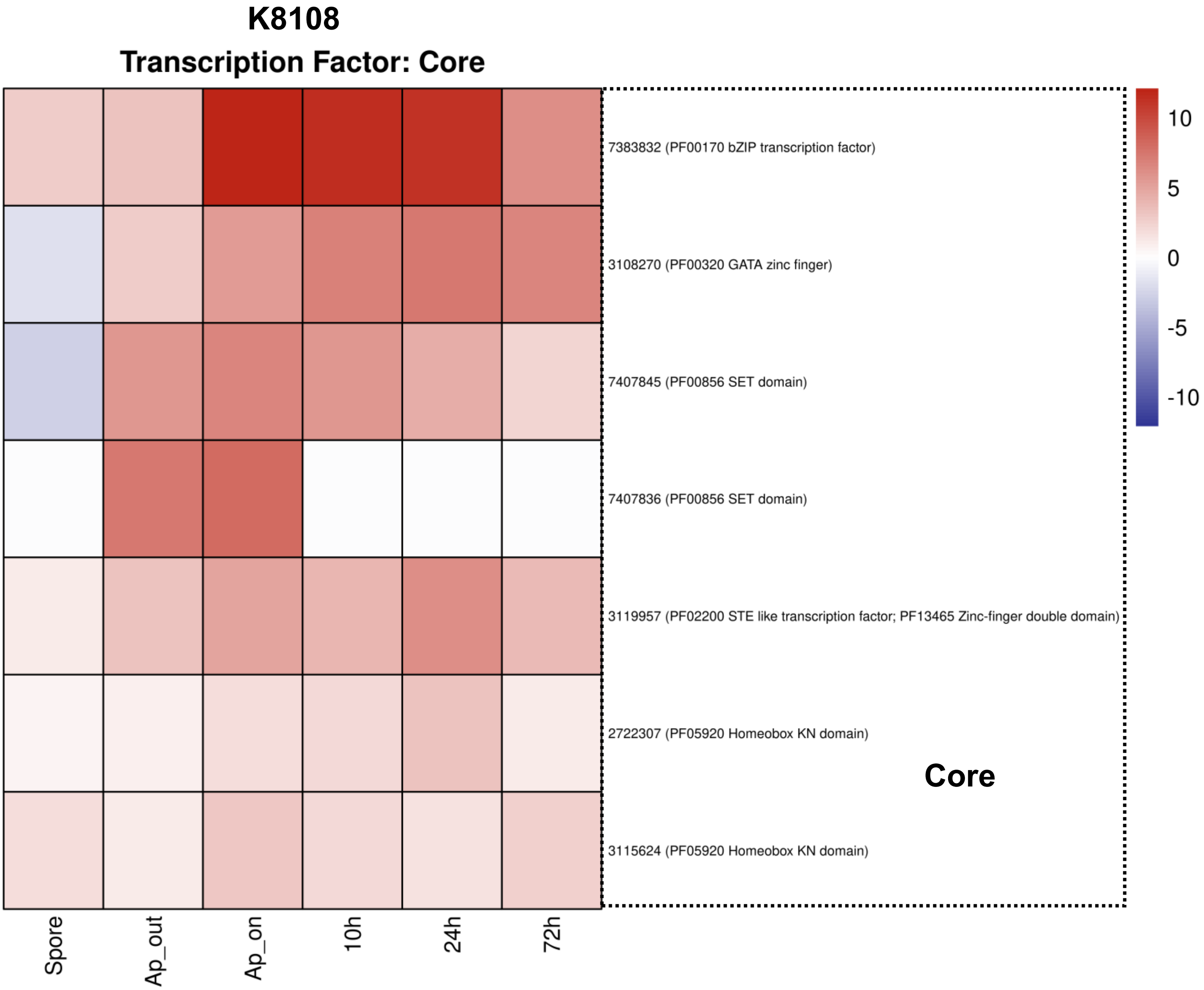


**Supplementary Figure S1-H**
